# MIRIAD: a Multiplex Immunoassay for Rodents Infectious and Animal Diseases

**DOI:** 10.1101/2025.10.24.684221

**Authors:** Ferdinando Scavizzi, Elisa Vandenkoornhuyse, Marcello Raspa, Clelia Buccheri, Christophe Audebert, Rémi Malbec

**Author notes:** Correspondence : Ferdinando Scavizzi,; Rémi Malbec.

## Abstract

Health monitoring of laboratory rodents is essential for ensuring animal welfare, meeting regulatory requirements for zoonosis control, and maintaining the integrity and reproducibility of scientific studies. Routine serological screening, conducted in accordance with FELASA guidelines, plays a critical role in the detection of infectious agents. While conventional serological assays typically target individual pathogens, existing multiplex platforms often require specialized equipment and complex protocols, limiting their routine use within animal facilities.

The MIRIAD® assay is a multiplex ELISA-based platform developed for in-house serological surveillance of rodents. It offers a user-friendly, reliable, and cost-effective alternative to outsourced testing. In this study, we evaluated the performance and usability of the MIRIAD® assay by analyzing serum samples from mice (n□=□68) and rats (n□= 27). Results were compared to two commercially available single-target ELISAs, with 94 % to 100 % concordance observed across all tested pathogens. Assay compatibility with dried blood spot (DBS) cards was also assessed, demonstrating reliable detection after storage and elution, and supporting the potential for reduced sample volumes and simplified handling.

These findings highlight MIRIAD® as a practical tool for routine health monitoring in laboratory animal facilities. Its ease of use, minimal sample requirements, and compatibility with DBS technology can facilitate more frequent testing, support early detection of pathogens, and promote adherence to the 3Rs principles through refinement of monitoring practices.

## 1. Introduction

Maintaining Specific Pathogen-Free (SPF) or Specific and Opportunistic Pathogen-Free (SOPF) status is critical for ensuring animal welfare, minimizing zoonotic risks, and preserving the reproducibility of biomedical research. The presence of undetected pathogens can introduce significant biological variability, leading to compromised data integrity. Routine health monitoring, guided by FELASA recommendations, is therefore essential and typically involves serology, molecular diagnostics (PCR/RT-PCR), microbial culture, and microscopic examination (Mähler Convenor *et al*., 2014; Livingston and Riley, 2003).

Recent innovations such as environmental PCR (ePCR) and exhaust air dust (EAD) sampling provide non-invasive alternatives to sentinel programs (Mähler Convenor *et al*., 2014; Livingston and Riley, 2003). However, serology remains indispensable, particularly for detecting persistent or low-shedding infections like parvoviruses, and for confirming PCR findings that may yield false positives due to high analytical sensitivity (Sykes *et al*, 2022; Buchheister and Bleich, 2021). Serological assays detect pathogen-specific antibodies in serum or dried blood spot (DBS) samples, offering a window into past exposure.

Multiplex serological platforms, including bead-based assays and MFIA™, offer high-throughput testing from minimal sample volumes (Höfler *et al*., 2014; Khan *et al*., 2005; Christopher-Hennings *et al*., 2013; Henderson *et al*., 2013; Brochot *et al*., 2022). Despite their advantages, these systems often require costly, specialized instrumentation and technical expertise, prompting many institutions to outsource testing. Outsourcing, while convenient, introduces delays, limits scheduling flexibility, and reduces control over data handling and diagnostic workflows. These constraints can hinder timely outbreak responses and adaptive colony management.

In-house diagnostic solutions that are cost-effective, flexible, and easy to implement could greatly enhance health surveillance capabilities. Ideally, such platforms would support multiplex testing using standard laboratory equipment and simplify logistics by eliminating the need for multiple reference shipment and storage.

The MIRIAD® assay is a multiplex ELISA-based platform designed for on-site serological monitoring of rodents. It enables simultaneous detection of antibodies against multiple pathogens in a user-friendly and low-cost format. This study assesses the performance, usability, and diagnostic concordance of the MIRIAD® assay compared to established single-target ELISAs. Additionally, we evaluate its compatibility with DBS cards to support alternative sample collection and storage methods. Together, these findings aim to determine the suitability of MIRIAD® as an in-house tool for FELASA-compliant health monitoring in laboratory animal facilities.

## 2. Results and discussion

A total of 22 mouse and 5 rat serum samples were analyzed using the MIRIAD® Annual multiplex ELISA kit and 46 mouse and 22 rat serum samples were analyzed using the MIRIAD® Quarterly multiplex ELISA kit. All rat samples were negative across all serological targets. In contrast, 37 out of 68 mouse samples (54%) tested positive for murine norovirus (MNV). Additional findings included positivity for murine hepatitis virus (MHV; 10%), mouse parvovirus (MPV; 15%), Theiler’s murine encephalomyelitis virus (TMEV; 9%), and one out of 22 mouse samples tested positive for Clostridium piliforme (Tyzzer’s disease). Multi-pathogen seropositivity was observed in several cases: as five samples tested positive for MNV, MHV, TMEV, and MPV (Table 1). These results are consistent with previous epidemiological reports highlighting the dominance of MNV as a prevalent pathogen in laboratory mice. A European study by Pritchett-Corning et al. (2009) found a 24% seroprevalence for MNV—significantly higher than MPV (3.64%) and MHV (3.25%) (Pritchett-Corning and Cosentino, 2009). The elevated pathogenic prevalence and the presence of TMEV and TD antibodies in this cohort, though less common, reflects the historical disease burden of the facilities selected for this specific reason for this study.

**Table 1:** Comparison of results obtained between the MIRIAD® kit and Charles River’s ELISA (n= 22 annual + 46 quarterly mouse sera) and Biotech’s ELISA (n=5 annual + 22 quarterly rat sera), and between raw sera or sera collected on transport card (n= 15 mouse sera + 3 rat sera).

|  | **MIRIAD** |  | **Charles River** |  |  |
| --- | --- | --- | --- | --- | --- |
| **Mouse targets** | **N Positives** | **% Positives** | **N Positives** | **% Positives** | **% Concordance** |
| **MNV** | 37/68 | 54 % | 37/68 | 54 % | 94 % (64/68) |
| **MHV** | 7/68 | 10 % | 11/68 | 16 % | 94 % |
| **MPV** | 10/68 | 15 % | 10/68 | 15 % | 100 % |
| **MVM** | 0/68 | 0 % | 0/68 | 0 % | 100 % |
| **TMEV** | 6/68 | 9 % | 7/68 | 10 % | 99 % |
| **EDIM** | 0/68 | 0 % | 0/68 | 0 % | 100 % |
| **ECTV** | 0/22 | 0 % | 0/22 | 0 % | 100 % |
| **LCMV** | 0/22 | 0 % | 0/22 | 0 % | 100 % |
| **KV** | 0/22 | 0 % | 0/22 | 0 % | 100 % |
| **MCMV** | 0/22 | 0 % | 0/22 | 0 % | 100 % |
| **POLY** | 0/22 | 0 % | 0/22 | 0 % | 100 % |
| **FL** | 0/22 | 0 % | 0/22 | 0 % | 100 % |
| **K87** | 0/22 | 0 % | 0/22 | 0 % | 100 % |
| **REO** | 0/22 | 0 % | 0/22 | 0 % | 100 % |
| **SEV** | 0/22 | 0 % | 0/22 | 0 % | 100 % |
| **HTNV** | 0/22 | 0 % | 0/22 | 0 % | 100 % |
| **CARB** | 0/22 | 0 % | 0/22 | 0 % | 100 % |
| **ECUN** | 0/22 | 0 % | 0/22 | 0 % | 100 % |
| **TD** | 1/22 | 5 % | 1/22 | 5 % | 100 % |
| **MPUL** | 0/22 | 0 % | 0/22 | 0 % | 100 % |
| **PVM** | 0/22 | 0 % | 0/22 | 0 % | 100 % |
| **Rat targets** | 0/27 | 0 % | 0 | 0 % | 100 % |
| **Mouse targets** | **Sera → MIRIAD** |  | **Transport card → MIRIAD** |  |  |
| **MNV** | 5/15 | 34 % | 5/15 | 34 % | 100% |
| **MHV** | 3/15 | 20 % | 4/15 | 27 % | 93 % |
| **Rat targets** | 0/3 | 0 % | 0/3 | 0 % | 100 % |

All positive results obtained with the MIRIAD® multiplex assay were confirmed using corresponding commercial simplex ELISAs (Charles River for mouse samples, Biotech for rats), yielding 94% concordance for MNV (with 33 samples positives for both assays) and MHV and almost 100% concordance for the rest of the targets, of lower prevalence. Minor discrepancies were expected, as different commercial assays may differ in their targeted antigenic domains, production methodologies, and immunoassay formats and specificities—factors that can collectively impact epitope availability and reactivity. In such cases, confirmatory testing, combined with an understanding of the facility’s historical disease burden, can aid in establishing the final diagnosis.

To assess the quantitative consistency of the MIRIAD® assay, spot signal intensity (arbitrary units) was compared to normalized net absorbance from Charles River ELISAs for the four most commonly detected pathogens. Pearson correlation coefficients were R=0.65 for MNV, R=0.56 for MHV, R=0.85 for MPV and R= 0.61 for TMEV (supplementary Fig. 1).

To evaluate compatibility with dried serum formats, 15 mouse and 3 rat sera were applied to transport cards and stored at 4□°C. Given the potential loss of antibodies during the elution process, we chose to incubate the eluates at a 1:50 dilution using the MIRIAD® quarterly and annual kits. For comparison, fresh sera were tested at a 1:100 dilution. A strong correlation in signal intensity was observed between the two sample types, with Pearson correlation coefficients of R = 0.99 for MNV and R = 0.75 for MHV (Fig. 1). Using the transport cards, 4 out of 15 mouse samples tested positive for MNV and 4 out of 15 for MHV.

**Figure 2:**
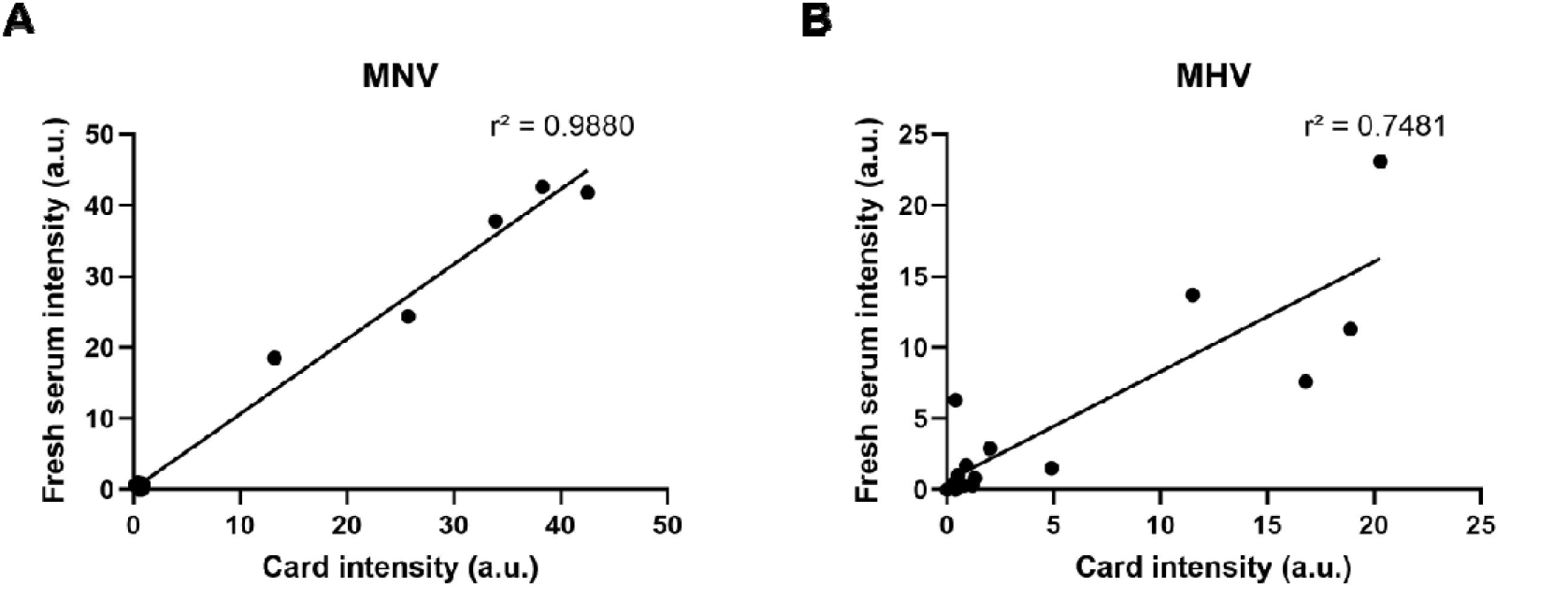
Correlation of spot intensity in arbitrary units (a.u.) for MNV (A) and MHV (B)targets between 100-fold dilution of fresh sera and 50-fold dilution of cards eluate using the MIRIAD® assay.

## 3. Discussion

This study demonstrates the reliability and operational value of the MIRIAD® multiplex ELISA for in-house health monitoring of rodents. The assay provided elevated concordance with commercial simplex tests and was compatible with dried serum formats. Its ease of use, minimal sample volume requirement, and capacity for multi-target screening make it suitable for routine implementation within animal facilities.

This approach clearly improves the respect of the 3Rs since only a smaller volume of serum is required allowing a less invasive and harmful blood sampling compared to single target ELISAs. Noteworthy, it makes possible to use, as few as 15 microliters, of dried blood to detect up to 26 pathogens.

Follow-up studies including additional pathogens and a larger cohort will further define the assay’s diagnostic breadth and utility.

## 5. Methods

Whole blood was collected from n□=□68 mice and n□=□27 rats housed in animal facilities of several Italian Research Centers and Universities. Small volume of blood (∼50 µl) were centrifuged to isolate serum, which was stored at −20□°C until analysis.

MIRIAD® assay protocol requires just 2 µl of serum for each multiplex-well (versus 10 µl of serum for each single-target well for standard commercial simplex assays). Thus, a less invasive and painful blood draw from facial vein or retro-orbital venous plexus is sufficient to this aim.

Serological screening was performed using commercial ELISA kits: a simplex ELISA from Charles River (Wilmington, MA) for mouse samples and from Biotech Trading Partners (Encinitas, CA, USA) for rat samples. In addition, all samples were tested using the multiplex MIRIAD® Annual or Quarterly kits (GD Biotech, Loos, France), following the manufacturer’s protocol.

The MIRIAD® assay is a miniaturized, multiplex ELISA platform in which each well is spotted with 26 pathogen-specific antigens (annual version) or 14 pathogen-specific antigens (quarterly version), enabling parallel detection of IgG antibodies in serum samples (supplementary Fig. 2). After incubation with serum, unbound material was removed by washing. An HRP-conjugated secondary antibody was then applied, followed by TMB substrate to generate an insoluble blue precipitate at antigen–antibody binding sites. The color intensity, corresponding to the relative abundance of specific IgGs, was quantified via bottom-well imaging using the MIRIAD® Reader (Malbec *et al*., 2020). Net spot intensities (Arbitrary Units, A.U.) were calculated, and serostatus was interpreted using the MIRIAD® Annual Mouse and Rat analysis software.

To assess compatibility with dried sample formats, serum transport cards (Innovative Diagnostics, Montpellier, France) were used. The serum cards are designed to collect and transport serum samples, indeed samples may be conserved on the card for up to 8 days and shipped at room temperature (21° C ± 5°C). After serum collection, 15 µl of blood was deposited on each card. Cards were allowed to dry for 10 minutes and serum was obtained by migration of whole blood, without centrifugation.

Serum from n□=□3 mice was applied to the cards and stored at 4□°C. After 5 days, samples were eluted according to the manufacturer’s instructions and tested at 1:50 and 1:100 dilutions using the MIRIAD® Annual kit. Results were compared to those obtained from matched fresh serum samples run under identical conditions.

**Sup. Figure 2:**
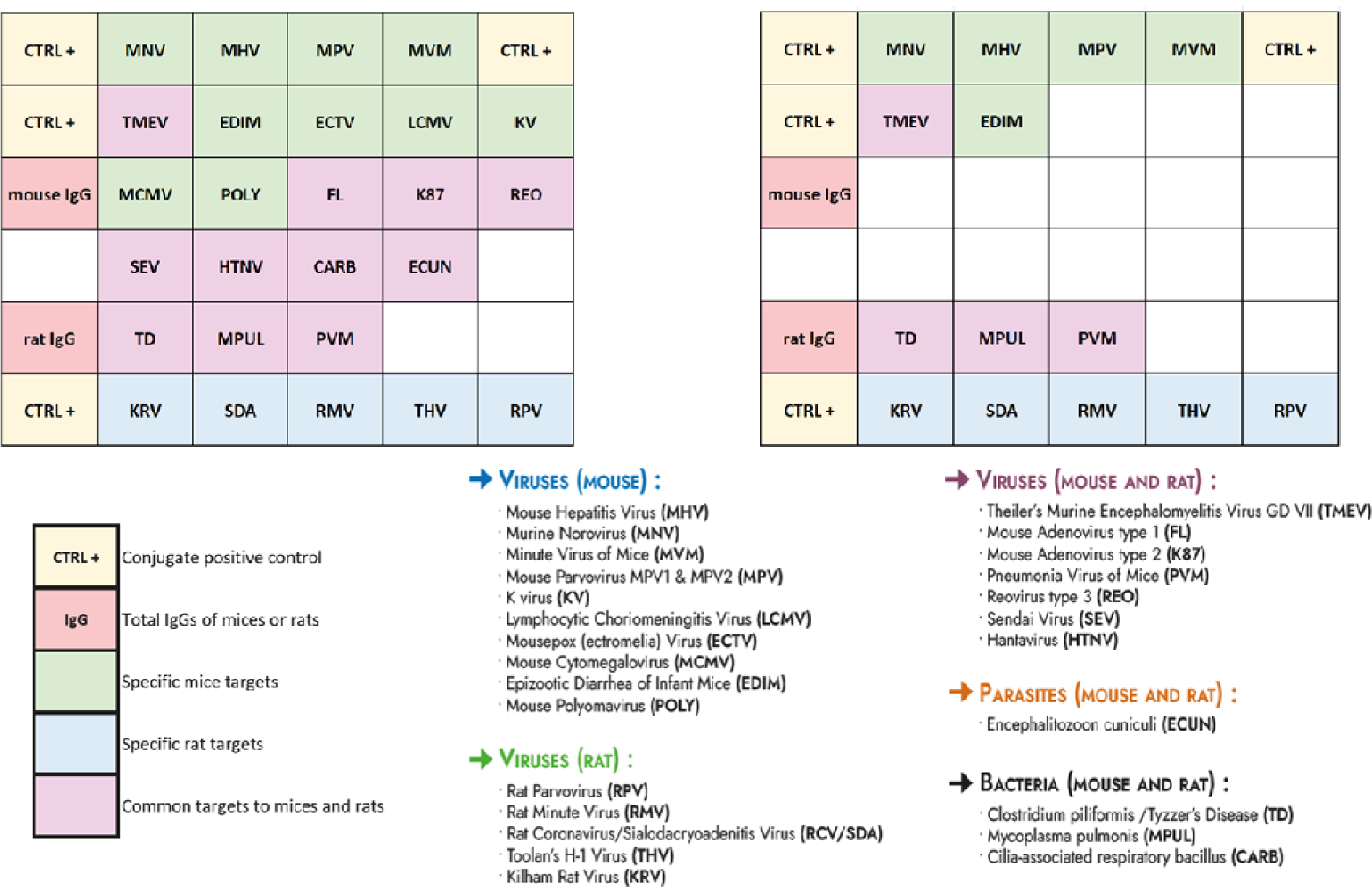
List of target pathogens in the MIRIAD® annual (A) and quarterly (B) kits and antigen array design at the bottom on each well (adapted from the MIRIAD ® instructions for use).

## 6. Statements and Declarations

### 6.3 Statements of Animal Ethics

The animal study protocol was approved by the Institutional Review Board (Organismo Preposto al Benessere degli Animali, OPBA) of the Institute of Biochemistry and Cell Biology— EMMA/Infrafrontier (Protocol number 0000079 of January 18, 2016) for studies involving animals.

## 7. Acknowledgements

Authors wish to thank Daniele Iannilli, Massimiliano Iannilli and Arsenio Armagno for excellent technical support.

## 8. Author contributions

F.S. conceptualized the study, designed the study, supported data collection, analyzed the data, drafted the manuscript and revised the manuscript.

C.B. and R.M ran the assays, collected and analyzed the data.

E.V. and C.A. revised the manuscript.

