## Supplemental Figure 1 for "MIRIAD: a Multiplex Immunoassay for Rodents Infectious and Animal Diseases"

### Sup. Figure 1

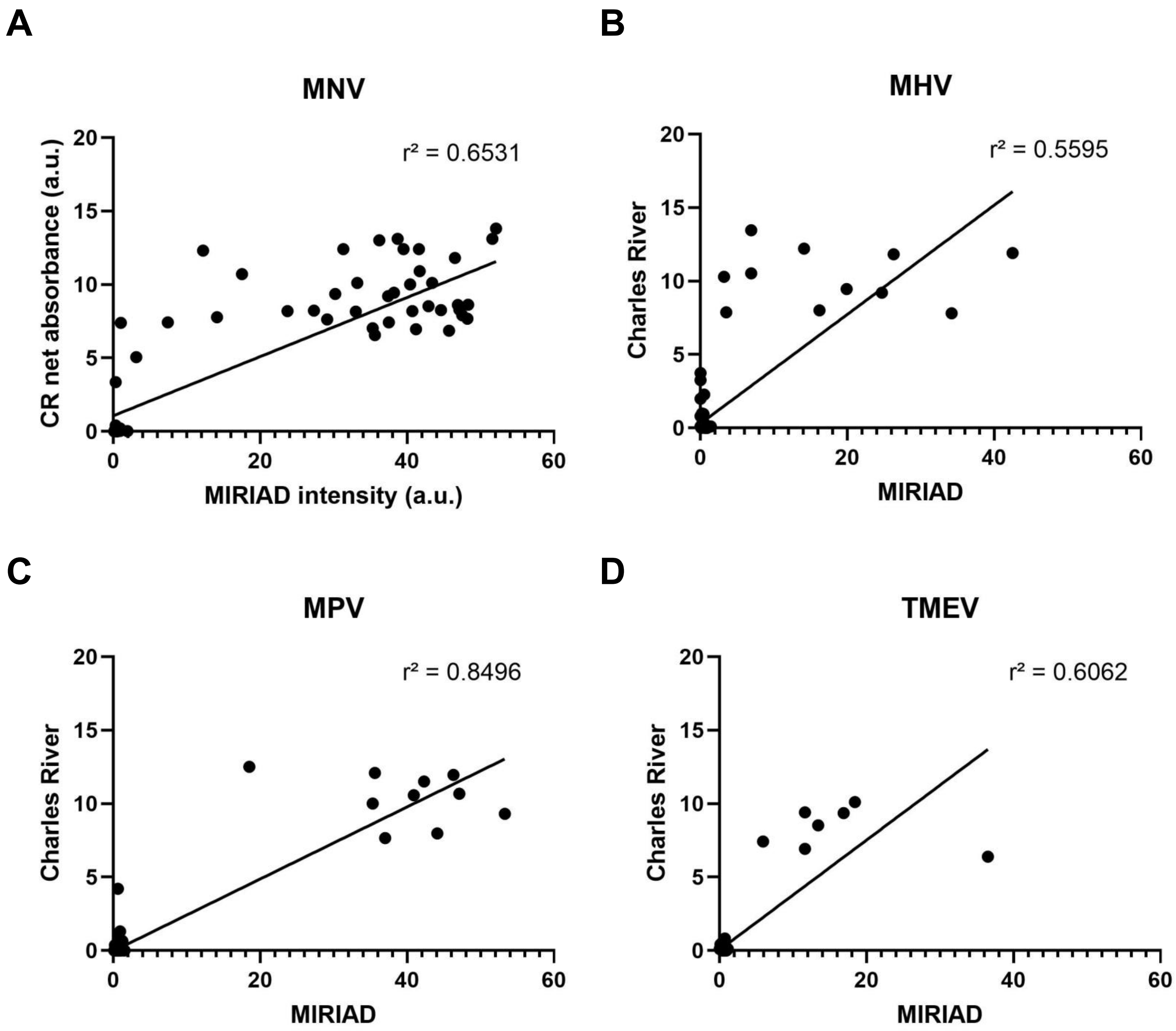


**Sup. Fig 1 : Charles River normalized net absorbance (a.u.) against MIRIAD spot intensity in arbitrary units (a.u.) for MNV (A), MHV (B), MPV (C), and TMEV (D) targets.**
