## Supplemental Figure 2 for "MIRIAD: a Multiplex Immunoassay for Rodents Infectious and Animal Diseases"

### Sup. Figure 2

**A**

| | | | | | |
|---|---|---|---|---|---|
| CTRL + | MNV | MHV | MPV | MVM | CTRL + |
| CTRL + | TMEV | EDIM | ECTV | LCMV | KV |
| mouse IgG | MCMV | POLY | FL | K87 | REO |
| | SEV | HTNV | CARB | ECUN | |
| rat IgG | TD | MPUL | PVM | | |
| CTRL + | KRV | SDA | RMV | THV | RPV |

**B**

| | | | | | |
|---|---|---|---|---|---|
| CTRL + | MNV | MHV | MPV | MVM | CTRL + |
| CTRL + | TMEV | EDIM | | | |
| mouse IgG | | | | | |
| rat IgG | TD | MPUL | PVM | | |
| CTRL + | KRV | SDA | RMV | THV | RPV |

| | |
|---|---|
| CTRL + | Conjugate positive control |
| IgG | Total IgGs of mices or rats |
| | Specific mice targets |
| | Specific rat targets |
| | Common targets to mices and rats |

→ **Viruses (mouse) :**

- Mouse Hepatitis Virus **(MHV)**
- Murine Norovirus **(MNV)**
- Minute Virus of Mice **(MVM)**
- Mouse Parvovirus MPV1 & MPV2 **(MPV)**
- K virus **(KV)**
- Lymphocytic Choriomeningitis Virus **(LCMV)**
- Mousepox (ectromelia) Virus **(ECTV)**
- Mouse Cytomegalovirus **(MCMV)**
- Epizootic Diarrhea of Infant Mice **(EDIM)**
- Mouse Polyomavirus **(POLY)**

→ **Viruses (rat) :**

- Rat Parvovirus **(RPV)**
- Rat Minute Virus **(RMV)**
- Rat Coronavirus/Sialodacryoadenitis Virus **(RCV/SDA)**
- Toolan's H-1 Virus **(THV)**
- Kilham Rat Virus **(KRV)**

→ **Viruses (mouse and rat) :**

- Theiler's Murine Encephalomyelitis Virus GD VII **(TMEV)**
- Mouse Adenovirus type 1 **(FL)**
- Mouse Adenovirus type 2 **(K87)**
- Pneumonia Virus of Mice **(PVM)**
- Reovirus type 3 **(REO)**
- Sendai Virus **(SEV)**
- Hantavirus **(HTNV)**

→ **Parasites (mouse and rat) :**

- Encephalitozoon cuniculi **(ECUN)**

→ **Bacteria (mouse and rat) :**

- Clostridium piliformis /Tyzzer's Disease **(TD)**
- Mycoplasma pulmonis **(MPUL)**
- Cilia-associated respiratory bacillus **(CARB)**

**Sup. Fig. 2 List of target pathogens in the MIRIAD® annual (A) and quarterly (B) kits and antigen array design at the bottom on each well (adapted from the MIRIAD ® instructions for use).**
